# Required marker properties for unbiased estimates of the genetic correlation between populations

**DOI:** 10.1101/301333

**Authors:** Yvonne C.J. Wientjes, Mario P.L. Calus, Pascal Duenk, Piter Bijma

**Affiliations:** Wageningen University & Research, Animal Breeding and Genomics, 6700 AH Wageningen, The Netherlands

**Author notes:** Author information: Yvonne Wientjes Wageningen University & Research Animal Breeding and Genomics P.O. box 338, 6700 AH Wageningen, the Netherlands.

**Keywords:** genetic correlation between populations, genomic relationships, marker-based relationships, multi-trait model

## Abstract

Populations generally differ in environmental and genetic factors, which can create differences in allele substitution effects between populations. Therefore, a single genotype may have different additive genetic values in different populations. The correlation between the two additive genetic values of a single genotype in both populations is known as the additive genetic correlation between populations and can differ from one. Our objective was to investigate whether differences in linkage disequilibrium (LD) and allele frequencies of markers and causal loci between populations affect bias of the estimated genetic correlation. We simulated two populations that were separated for 50 generations. Markers and causal loci were selected to either have similar or different allele frequencies in the two populations. Differences in consistency of LD between populations were obtained by using different marker density panels. Results showed that when the difference in allele frequencies of causal loci between populations was reflected by the markers, genetic correlations were only slightly underestimated using markers. This was even the case when LD patterns, measured by LD statistic *r*, were different between populations. When the difference in allele frequencies of causal loci between populations was not reflected by the markers, genetic correlations were severely underestimated. We conclude that for an unbiased estimate of the genetic correlation between populations, marker allele frequencies should reflect allele frequencies of causal loci so that marker-based relationships can accurately predict the relationships at causal loci, i.e. *E*(**G**_causal loci_|**G**_markers_) ≠ **G**_markers_. Differences in LD between populations have little effect on the estimated genetic correlation.

## INTRODUCTION

Alleles in different populations are often expressed in a different environment and a different genetic background. As a result of genotype by environment interaction and non-additive genetic effects, those differences result in different allele substitution effects between populations (Fisher 1918; Fisher 1930; Falconer 1952). In addition, the set of loci underlying a trait can differ between populations. Therefore, a single genotype may have different additive genetic values in different populations. For each population, the additive genetic value is the product of the genotype, measured as allele count at each locus, multiplied by the allele substitution effects for that population. The additive genetic correlation between two populations is the correlation between the two additive genetic values of a single genotype in both populations and may considerably differ from one.

Knowledge of the genetic correlation between populations helps to understand the differences and similarities between populations in genetic architecture of complex traits (De Candia *et al.* 2013; Brown *et al.* 2016). For both genomic prediction and genome-wide association studies, combining information from populations is an attractive approach to increase the prediction accuracy of estimated genetic values or the power to identify quantitative trait loci. This is especially the case when the number of individuals with genotypes and phenotypes in a population is limited. For both genomic prediction as well as genome-wide association studies, the genetic correlation between populations determines the added benefit of combining information from multiple populations (De Candia *et al.* 2013; Wientjes *et al.* 2015; Wientjes *et al.* 2016). Therefore, the genetic correlation between populations is an important parameter in human studies (e.g., De Candia *et al.* 2013; Yang *et al.* 2013), as well as in animal and plant breeding (e.g., Karoui *et al.* 2012; Lehermeier *et al.* 2015).

For estimating a genetic correlation between two populations, it is essential to know the relationships between individuals from the two populations. Traditionally, relationships between individuals are based on pedigree information, which is generally only available within population. The current availability of genome-wide marker panels has opened up new opportunities to estimate genetic correlations between populations of distantly related individuals, such as between breeds (e.g., Karoui *et al.* 2012; Carillier *et al.* 2014), lines (Huang *et al.* 2014), sub-populations (e.g., Lehermeier *et al.* 2015), or ethnicities (e.g., De Candia *et al.* 2013; Yang *et al.* 2013). Genetic correlations between populations can be estimated using methods based on genomic relationships (Karoui *et al.* 2012), random regression on genotypes (Sørensen *et al.* 2012; Krag *et al.* 2013), or summary statistics of genome-wide association studies (Bulik-Sullivan *et al.* 2015; Brown *et al.* 2016). Wientjes *et al.* (2017) showed that an unbiased estimate of the genetic correlation can be obtained from genomic relationships based on causal loci.

Because causal loci are generally unknown, genomic relationships have to be based on marker information. The strength and phase of linkage disequilibrium (LD) between causal loci and markers is different between populations in humans (Sawyer *et al.* 2005), livestock (e.g., Heifetz *et al.* 2005; Veroneze *et al.* 2013) and plants (Flint-Garcia *et al.* 2003; Lehermeier *et al.* 2014). Due to imperfect LD between causal loci and markers, not all genetic variance is explained by the markers which can distort the estimation of genetic correlations (Bulik-Sullivan *et al.* 2015; Gianola *et al.* 2015). However, in a simulation study where populations had different LD patterns, the genetic correlation between populations was accurately estimated based on marker information (Wientjes *et al.* 2015).

The objective of this study was to investigate whether differences in LD and allele frequencies of markers and causal loci between populations affect bias of the estimated genetic correlation. We simulated two populations that were separated for 50 generations using scenarios differing in consistency of LD and in allele frequencies of markers and causal loci between the populations. We used different marker-based relationship matrices to estimate the genetic correlation.

## MATERIALS AND METHODS

### Population structure

Two populations were simulated using QMSim software (Sargolzaei and Schenkel 2009). The simulations were set-up to have the following two characteristics; 1) the two populations should have different LD patterns, as measured by the LD statistic *r*, and 2) a large number of loci should segregate in the last generation of which a part (>200 000) has similar allele frequencies in both populations and another part (>200 000) different allele frequencies in both populations. We simulated a historical population for 212 generations. The first generation (generation −211) contained 300 individuals. In the following 100 generations (generation −211 – −112), population size gradually decreased to 50 individuals to create LD. From generation −111 to generation −12, population size gradually increased to 300 individuals and was kept constant for the next 10 generations (generation −11 - −2). In the last generation of the historical population (generation −1), population size increased to 1800 individuals.

The last generation of the historical population was randomly divided into two equally sized populations (A and B) of 900 individuals. In the next generation, the size of both populations was increased to 1800 individuals and was kept constant for the following 40 generations (generation 1-40). Those reasonably large population sizes limited the drift of allele frequencies. Number of offspring was set to 10 and selection was at random, so the number of selected offspring per individual approximately followed a Poisson distribution, as assumed in the Wright-Fisher model of genetic drift. In the last 10 generations (generation 41-50), population size decreased to 120 individuals in each population to increase the extent of LD in each population, and the number of offspring was set to 20. In the entire simulation, the male to female ratio was 1:5, generations were not overlapping and mating was at random. All individuals from the last generation (2000) were used for the analyses.

### Genome size

A genome of 10 chromosomes of one Morgan each was simulated. This genome size was a balance between the computational effort of the analyses and the variation in relationships between family members. By using fewer chromosomes, computational effort reduced, but variation in relationships around their expectation based on the pedigree would have been inflated (Hill 1993). Each chromosome contained 300 000 randomly spaced loci, with a recurrent mutation rate of 0.00005 in the historical population. In the last generation of the historical population, segregating loci were selected and mutation was stopped. The chosen population size and mutation rate resulted in a U-shaped allele frequency distribution of loci in the two populations, as commonly found in real populations.

In the last generation (generation 50), markers and 2000 causal loci were selected from all segregating loci. Three marker panels were constructed: a High Density Panel (HDP) with 200 000 markers, a Low Density Panel (LDP) with 20 000 markers, and a Very Low Density Panel (VLDP) with 2000 markers. Each of the smaller marker panels was a subset from the larger marker panels. The different marker densities were used to represent differences in consistency of LD between populations, since consistency in LD decreases when genomic distance between markers and causal loci increases (De Roos et al. 2008).

Markers and causal loci were selected to either have similar or different allele frequencies in population A and B. For both approaches, three selection criteria were used; namely (1) the segregation in one or both populations, (2) the absolute difference in allele frequency between population A (*p_A_*) and population B (*p_B_*), and (3) the difference in variance explained by a locus between population A and B, when allele substitution effects would be the same in both populations. The last criterion was mainly effective for loci with a low allele frequency, since an apparently small difference in allele frequency can result in a relatively large difference in variance explained for those loci.

For selecting markers with similar allele frequencies in the two populations, loci had to (1) segregate in both populations, (2) |*p_A_* – *p_B_*| should be less than 0.14, and (3) |2*p_A_*(1–*p_A_*)–2*p_B_*(1–*p_B_*)|/[2*p̄_AB_*(1–*p̄_AB_*)] should be less than 2, where *p̄_AB_* was the average of *p_A_* and *p_B_*. For selecting markers with different allele frequencies in the two populations, (1) loci had to segregate in at least one population, (2) |*p_A_*–*p_B_*| should be more than 0.14, and (3) |2*p_A_*(1–*p_A_*) – 2*p_B_*(1–*p_B_*)/[2*p̄_AB_*(1–*p̄_AB_*)] should be more than 1. The cut-off values were chosen to either minimize or maximize the difference in allele frequencies between the populations, while ensuring that enough loci in each replicate met the criteria. We aimed to select marker panels with a uniform allele frequency distribution to reflect commercially available marker chips (Matsuzaki *et al.* 2004; Matukumalli *et al.* 2009; Ramos *et al.* 2009; Groenen *et al.* 2011). For this step, the loci that met the criteria were divided in 50 bins based on average allele frequency over the two populations (i.e., allele frequencies of bin 1 ranged from 0 – 0.02, of bin 2 from 0.02 – 0.04, etc.) and from each bin an equal number of loci was randomly selected. When the number of loci was too small in the two extreme bins (0.00 – 0.02, and 0.98 – 1.00), the bins were combined with the neighboring bin.

For selecting causal loci, the same criteria and cut-off values were used as for markers, with one exception. For the scenario where allele frequencies in the two populations were similar, causal loci did not have to segregate in both populations, since some causal loci are known to be at least partly population-specific (Kemper *et al.* 2015). As an additional criterion, causal loci could not already be selected as marker. Causal loci were randomly selected from all loci that met the criteria, and therefore their allele frequency pattern followed an approximate U-shaped distribution as expected for causal loci (Yang *et al.* 2010; Kemper and Goddard 2012).

### LD patterns and consistency of LD

The LD pattern and consistency in LD between the populations was investigated. Within each population and between all causal loci and markers less than 10 cM apart, the parameter *r* was calculated (Hill and Robertson 1968):

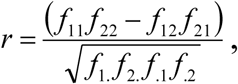

where *f_11_* is the haplotype frequency with allele 1 at the first locus and allele 1 at the second locus, *f_22_*, *f_12_* and *f_21_* are frequencies of the other possible haplotypes, *f_1_* and *f_2_*. are the frequencies of allele 1 and allele 2 at the first locus, and *f._1_* and *f._2_* are the frequencies of allele 1 and allele 2 at the second locus. The LD pattern within each population was represented by the average *r^2^* for intervals of 0.1 cM distance between the markers. The consistency of LD between the two populations was calculated as the correlation between *r* values of the two populations for intervals of 0.1 cM, following De Roos *et al.* (2008).

### Phenotypes

For each causal locus, allele substitution effects were sampled from a bivariate normal distribution, with mean 0, standard deviation 1, and a correlation between the populations of either 1, 0.8, 0.6, 0.4, 0.2 or 0. For each individual, its allele counts for the causal loci (coded as 0, 1, and 2) were multiplied by the corresponding allele substitution effects and results were summed over loci to calculate the additive genetic value (AGV) of the individual. The AGV were scaled to a mean of 0 and variance of 1 across all individuals. Since allele substitution effects were sampled independently from allele frequency, the correlation between AGV of population 1 and 2 (i.e., genetic correlation) was similar to the correlation between allele substitution effects (i.e., either 1, 0.8, 0.6, 0.4, 0.2 or 0). A normally-distributed environmental effect was sampled for each individual to obtain a heritability of 0.3 in each population. Phenotypes of all 2000 individuals in generation 50 were computed by summing the AGV and the environmental effects.

### Estimating the genetic correlation

The additive genetic correlation between populations was estimated using the following bivariate model:

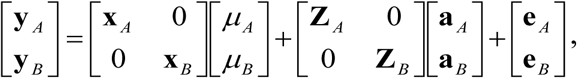

where **y**_*k*_ is a vector with phenotypes for population *k* (*k*= *A*, *B*), **x**_*k*_ is an incidence vector relating phenotypes to the mean in population *k* (*μ_k_*), **Z**_*k*_ is an incidence matrix relating phenotypes to estimated additive genetic values 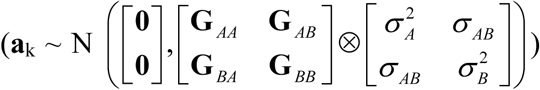 with ⊗ representing the Kronecker product function, and **e**_k_ are vectors with independent residual effects. Genetic and residual variances were estimated using REML. The first analyses were performed using ASReml software (Gilmour *et al.* 2015). For the scenarios analyzed later, we switched to MTG2 (Lee and van der Werf 2016) to reduce computation time. We verified that the estimated variance components were identical using both programs.

The genomic relationship matrix (**G**) between all individuals was calculated as (Wientjes *et al.* 2017):

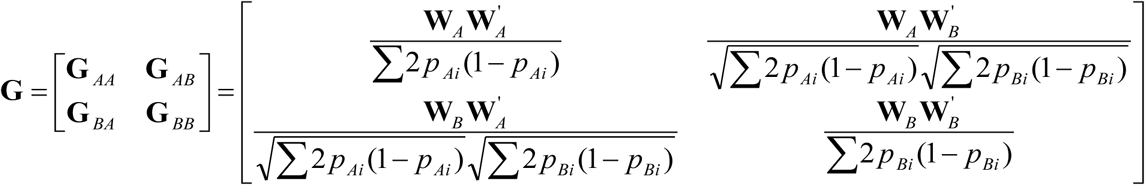

where **W**_*k*_ is a matrix with centered allele counts of all individuals from population *k*, and *p_ki_* is the allele frequency for locus *i* in population *k.* Centered allele counts were calculated as *g_ijk_* – 2*p_ik_*, where *g_ijk_* is the allele count of locus *i* for individual *j* from population *k*, coded as 0, 1 or 2. This **G** defines the relationships as standardized covariances between the genetic values of individuals (Wientjes *et al.* 2017). In all scenarios and in all 50 replicates, we calculated **G** using allele counts of 1) causal loci, 2) HDP markers, 3) LDP markers, or 4) VLDP markers.

The relationships at causal loci are the true relationships for that trait, that are approximated when using markers. Marker-based relationships are subject to sampling error, since markers are a subset of the genome. A way to account for this sampling error is by regressing **G** towards the pedigree relationship matrix (**A**) (Powell *et al.* 2010; Yang *et al.* 2010; Goddard *et al.* 2011), which is expected to reduce bias of estimated variance components (Yang *et al.* 2010). To investigate the effect of this regression, **G** matrices based on the three marker panels were regressed towards **A** and used for the scenarios with a correlation of 0.8 or 0.4.

Before regressing **G** towards **A**, the inbreeding level of each within-population block in **G** was rescaled to the inbreeding level in **A**, following (Powell *et al.* 2010):

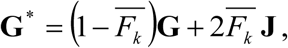

where *F_k_* is the average inbreeding coefficient of all individuals of population *k* based on the pedigree, and **J** is a matrix of ones. The rescaled **G**^*^ was regressed towards **A** following (Yang *et al.* 2010; Goddard *et al.* 2011):

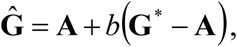

with

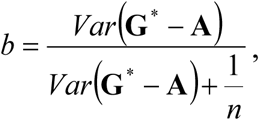

where *n* is the number of markers. To set-up **A**, the pedigree of the last 10 generations was used, so that between-population **A** relationships were zero. The regression was done separately within each population per bin of pedigree relationships (<0.10, 0.10-0.25, 0.250.50, >>0.5) and between populations, since regression coefficients are higher for higher pedigree relationships (Veerkamp *et al.* 2011; Wientjes *et al.* 2013). For the diagonal elements, only the inbreeding coefficients were regressed (Yang *et al.* 2010). Regression coefficients were all close to one for higher marker density panels (>0.99 for HDP and >0.97 for LDP). For VLDP markers, regression coefficients were lower; ∼0.84 for between-population relationships, ∼0.89, ∼0.91, ∼0.94 and ∼0.96 for the four bins of within-population relationships, and ∼0.93 for inbreeding coefficients.

### Data availability

Supplemental Material, File S1, is available at FigShare. This file contains the input file used for QMSim, the Fortran-programs to select markers and causal loci for the different scenarios, the Fortran-program to simulate phenotypes and the seeds for the different programs in each of the replicates.

## RESULTS

### Characteristics of simulations

The criteria for selecting markers and causal loci resulted in clear differences between the scenarios with similar and different allele frequencies in the two populations (Figure 1). As intended, the allele frequency distribution was uniform for markers and U-shaped for causal loci (not shown). Therefore, the percentage of causal loci with a minor allele frequency below 0.05 was higher (on average 33% in each population) than the percentage of markers with a minor allele frequency below 0.05 (on average only 15% in each population). The decay of LD was similar in both populations (Figure 2), with a strong decay of LD at increasing distances between the loci at the 0 – 2 cM interval. The consistency of LD phase decreased rapidly at short distances (0 – 5 cM), and fluctuated around zero at distances larger than 5 cM.

**Figure 1.**
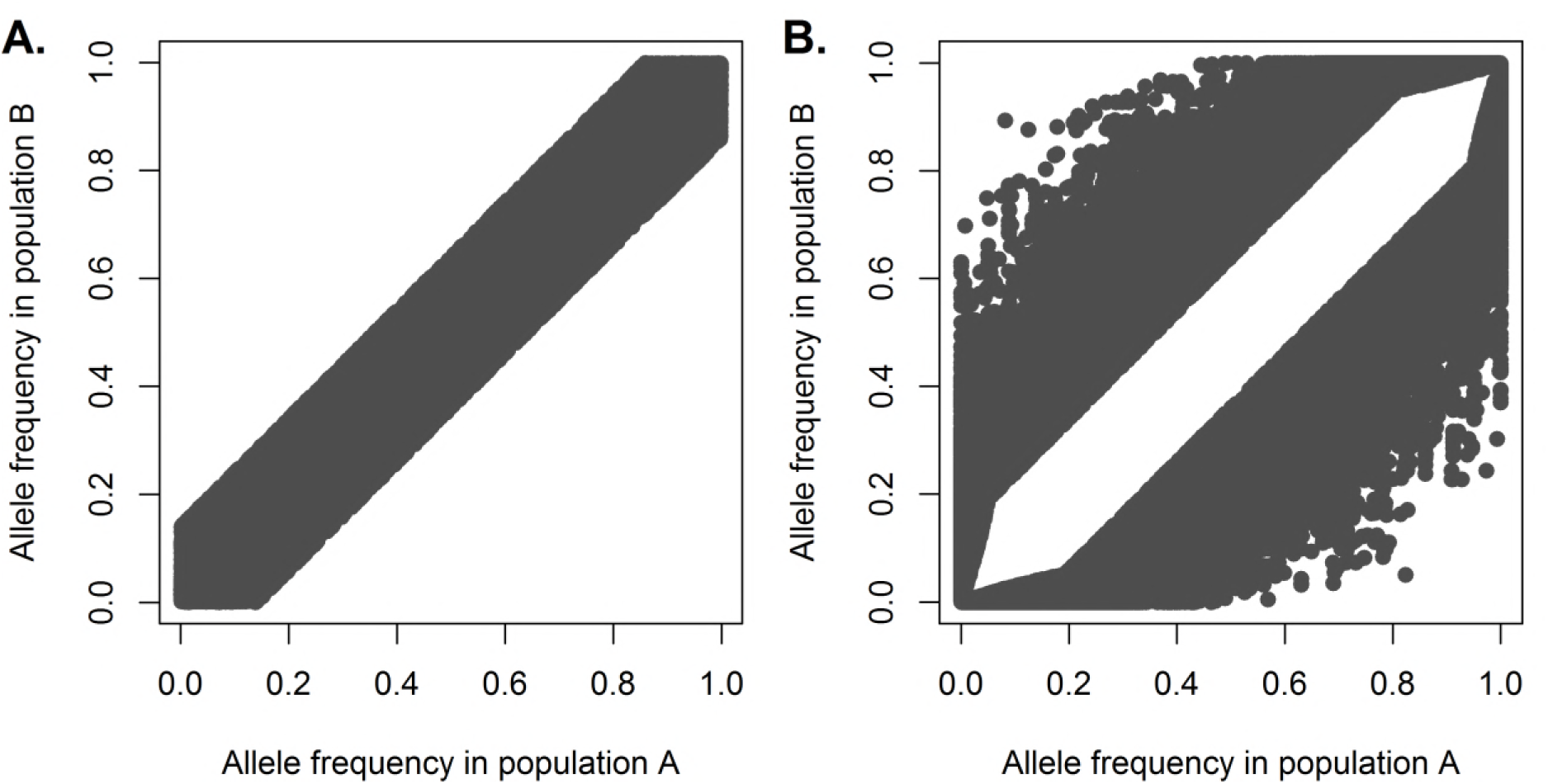
- Allele frequencies of markers for two populations using two selection approaches. For one random replicate, allele frequencies of markers from both populations are plotted against each other when markers are selected to have (**A.**) similar allele frequencies in the two populations, or (**B.**) different allele frequencies in the two populations.

**Figure 2.**
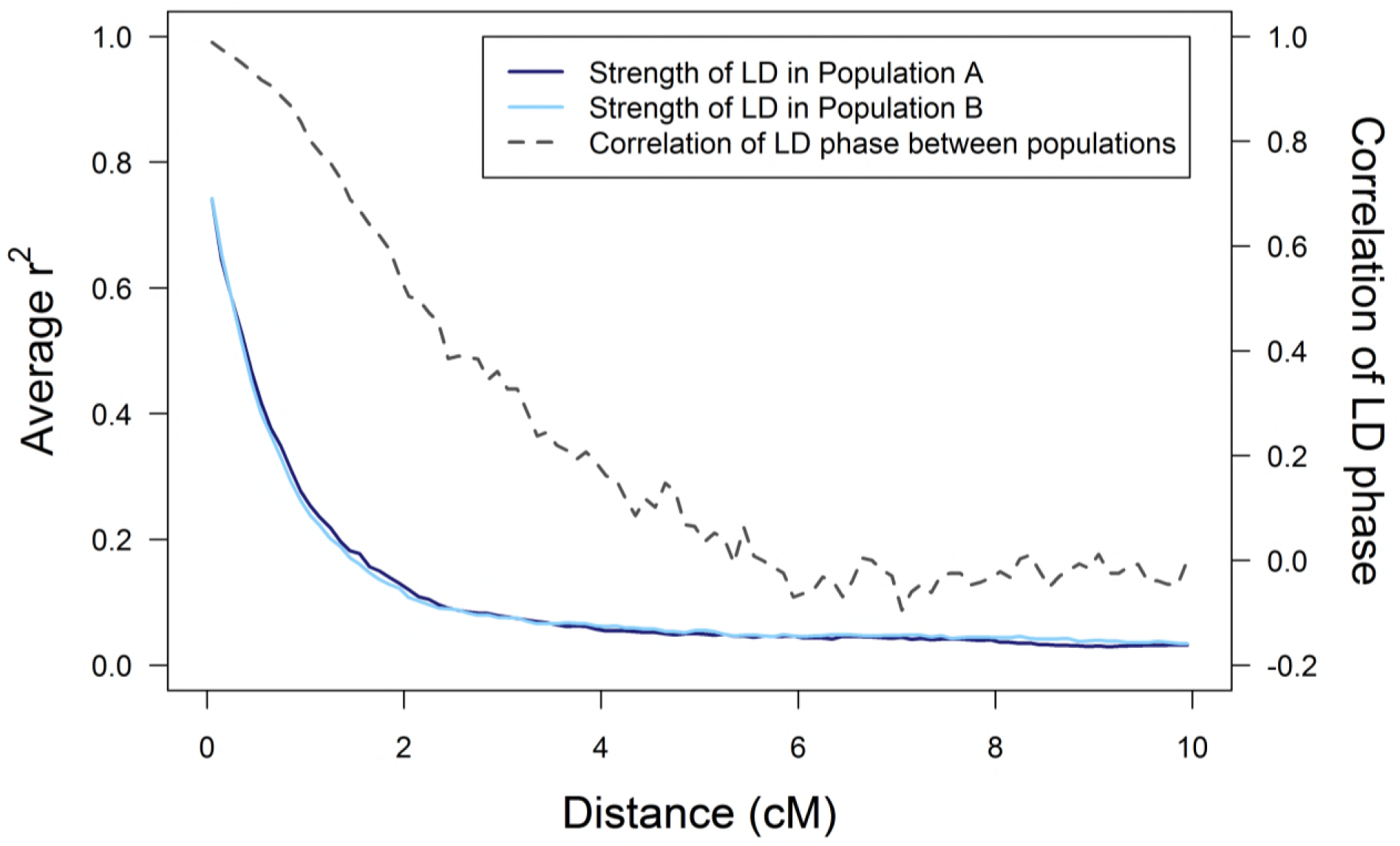
- LD pattern in two populations and correlation of LD phase between the populations. The average LD (*r^2^*) between causal loci and markers for both populations, and the correlation of LD-phase (correlation of *r*) between the populations, as a function of distance between causal loci and markers for one random replicate.

### Proportion of variance explained

The proportion of the phenotypic variance explained by the markers, known as the genomic heritability (De los Campos *et al.* 2015), was close to the simulated heritability for all scenarios (not shown). This implies that genetic variances were accurately estimated using all three marker panels.

### Estimated genetic correlation

With relationships based on causal loci, all estimated genetic correlations were unbiased, irrespective of whether causal loci had similar or different allele frequencies in the two populations (Figure 3). This was also expected based on previous results (Wientjes *et al.* 2017).

**Figure 3.**
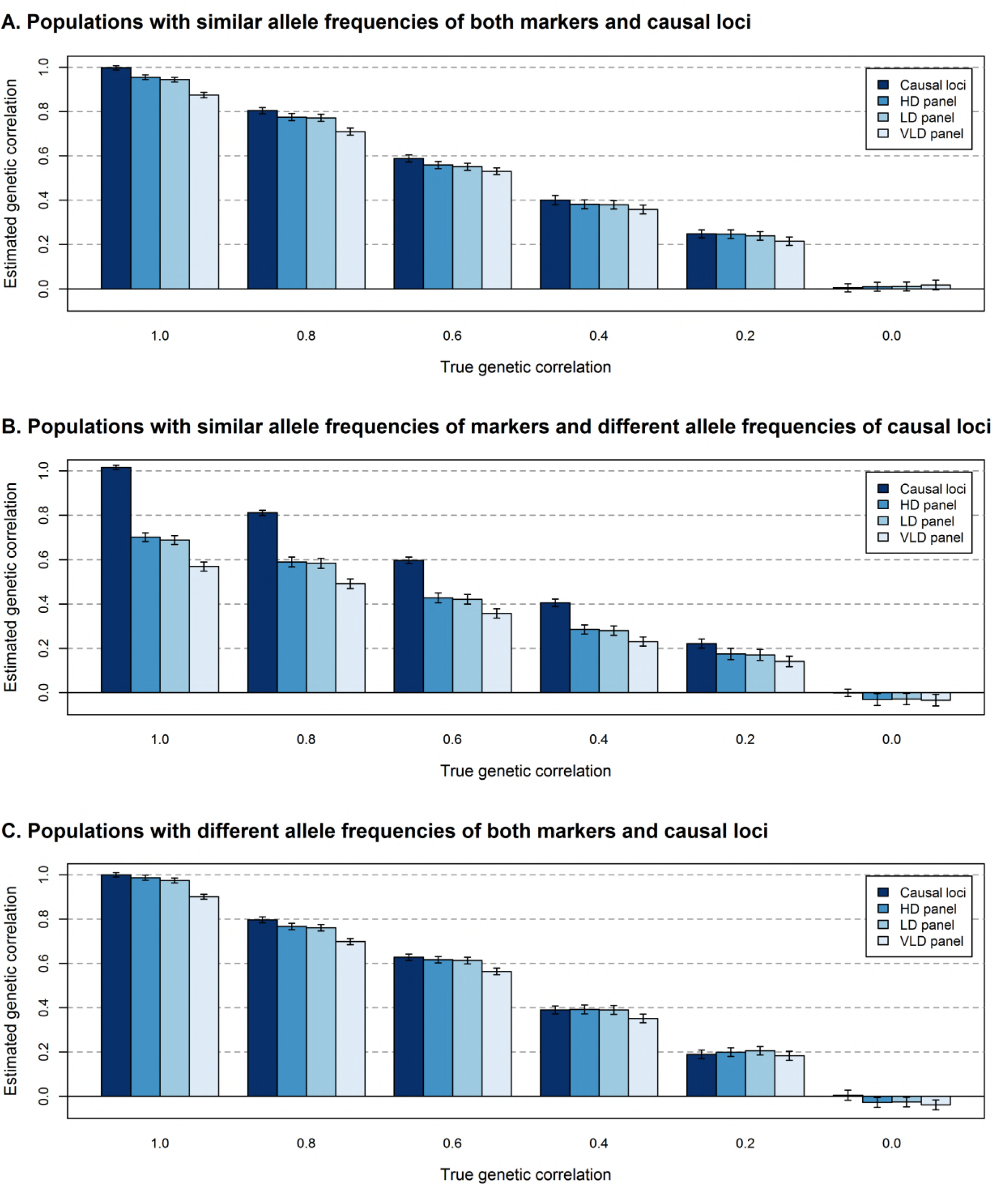
- Estimated genetic correlations between populations without regressing the genomic relationship matrix. The average estimated genetic correlation (± standard error) at different simulated genetic correlations for the scenario where (**A.**) markers and causal loci have similar allele frequencies in the two populations, (**B.**) markers have similar and causal loci different allele frequencies in the two populations, or (**C.**) markers and causal loci have different allele frequencies in the two populations, when the genomic relationship matrix is either based on the genotypes of causal loci (2000), HDP (200 000), LDP (20 000), or VLDP (2000) markers without regression towards the pedigree relationship matrix. Standard errors were calculated as the standard deviation over replicates divided by the square root of the number of replicates.

With relationships based on markers, all estimated genetic correlations were biased. When marker-based relationships were not regressed towards the pedigree relationships, genetic correlations were only slightly underestimated when the difference in allele frequencies of causal loci between populations was reflected by the markers, i.e., when markers and causal loci both had similar or different allele frequencies in the two populations (Figure 3A and 3C; ∼2.5% for HDP, ∼3% for LDP, and ∼11% for VLDP). The genetic correlation was much more severely underestimated when the difference in allele frequencies of causal loci between populations was not reflected by the markers (Figure 3B; ∼28% for HDP, ∼30% for LDP, and ∼41% for VLDP).

Across all scenarios, regressing **G** towards the pedigree relationship matrix only had a small effect on the estimated genetic correlation (Figure 4). At a high marker density, regressing **G** lowered the estimated genetic correlation. Therefore, the underestimation for HDP and LDP markers increased from ∼4% to ∼9% when the difference in allele frequencies of causal loci between populations was reflected by the markers, and from ∼28% to ∼32% when the difference in allele frequencies of causal loci between populations was not reflected by the markers. In contrast, regressing **G** resulted in higher estimated genetic correlations at low marker density. For VLDP markers, the underestimation decreased from ∼12% to ∼8% when the difference in allele frequencies of causal loci between populations was reflected by the markers, and from ∼41% to ∼38% when the difference in allele frequencies of causal loci between populations was not reflected by the markers. Thus, regressing **G** was only beneficial for estimating the genetic correlation between populations when the marker density was low.

**Figure 4.**
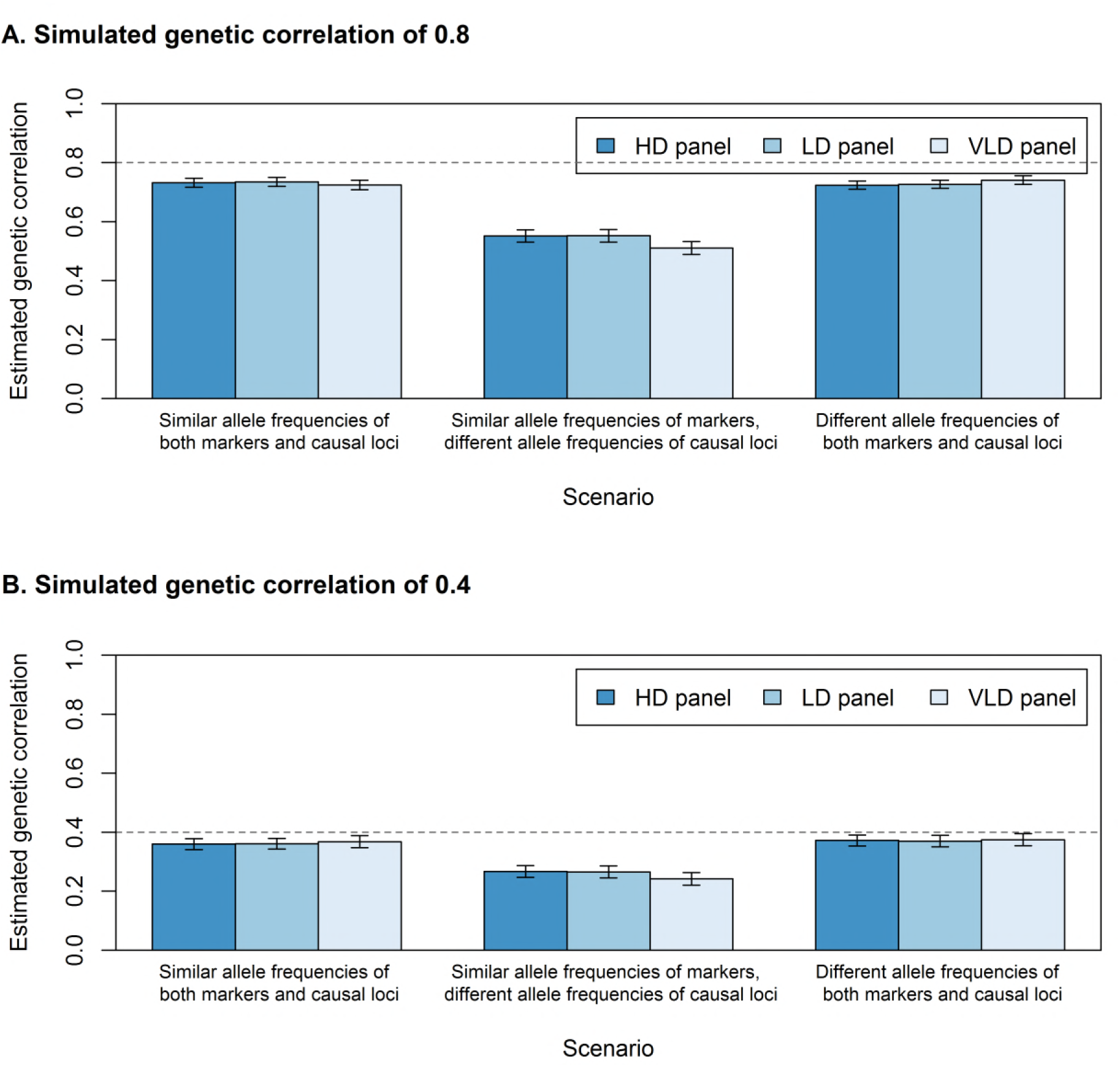
- Estimated genetic correlations between populations with regression of the genomic relationship matrix. The average estimated genetic correlation (± standard error) at a simulated genetic correlation of (**A.**) 0.8 or (**B.**) 0.4 for the three scenarios with HDP (200 000), LDP (20 000), or VLDP (2000) markers and regression of **G** towards the pedigree relationship matrix. Standard errors were calculated as the standard deviation over replicates divided by the square root of the number of replicates.

Standard errors across replicates for the estimated genetic correlation were generally small for all scenarios (∼0.02), and tended to be slightly larger for lower true genetic correlations. Moreover, standard errors were slightly larger when the difference in allele frequencies of causal loci between populations was not reflected by the markers (Figure 3B versus Figure 3A and 3C). Regression of **G** towards the pedigree relationship matrix had no effect on the standard error.

### Genomic relationships

Genetic variance estimates are biased when the regression of true relationships on marker-based relationships is not equal to one (Goddard *et al.* 2011). We investigated whether this could explain the underestimation of the genetic correlation by considering the genomic relationships at the causal loci as the true relationships for that trait. In Figure 5 and 6, we plotted the relationships at the causal loci versus the unregressed relationships at the markers for one of the replicates. The regression coefficients for within-population genomic relationships were close to one, and were only slightly lower when causal loci had different allele frequencies (Figure 6) compared to similar allele frequencies (Figure 5) in the two populations. This means that the within-population relationships at the markers can quite accurately predict the relationships at the causal loci.

**Figure 5.**
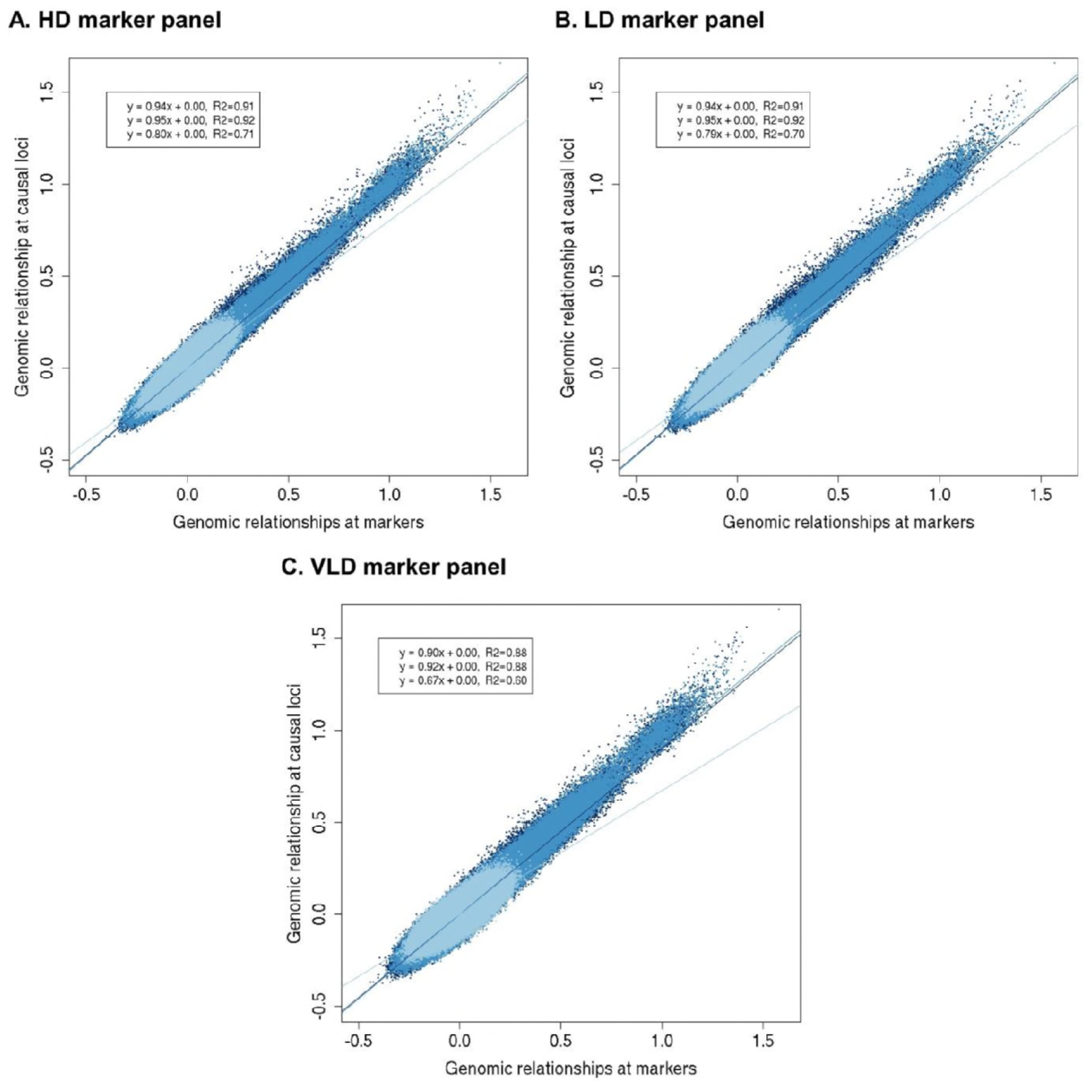
- Genomic relationships at causal loci versus markers when causal loci have similar allele frequencies in the two populations. The genomic relationships at the causal loci versus the genomic relationships based on (**A.**) HDP (200 000) markers, (**B.**) LDP (20 000) markers, or (**C.**) VLDP (2000) markers, when markers and causal loci have similar allele frequencies in the two populations for one replicate. Relationships in population A are represented in dark blue (equation 1 of regression line and correlation), relationships in population B are represented in medium blue (equation 2 of regression line and correlation), and relationships between population A and B are represented in light blue (equation 3 of regression line and correlation).

**Figure 6.**
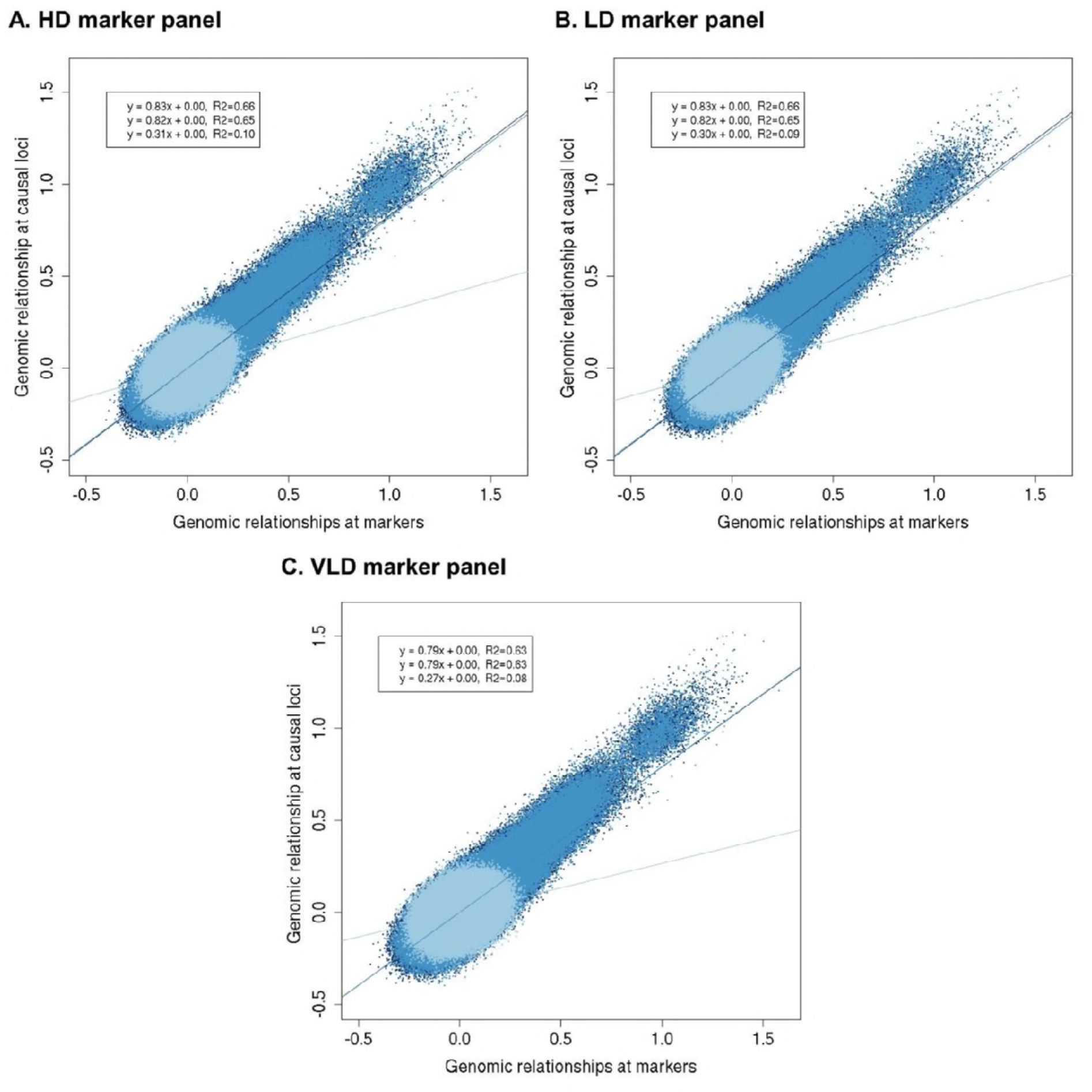
- Genomic relationships at causal loci versus markers when causal loci have different allele frequencies in the two populations. The genomic relationships at the causal loci versus the genomic relationships based on the (**A.**) HDP (200 000) markers, (**B.**) LDP (20 000) markers, or (**C.**) VLDP (2000) markers, when markers have similar and causal loci different allele frequencies in the two populations for one replicate. Relationships in population A are represented in dark blue (equation 1 of regression line and correlation), relationships in population B are represented in medium blue (equation 2 of regression line and correlation), and relationships between population A and B are represented in light blue (equation 3 of regression line and correlation).

Regression coefficients of between-population relationships deviated more from one, especially at low marker density. When the difference in allele frequencies of causal loci between populations was reflected by the markers, the regression coefficients were ∼0.8 for HDP and LDP, and 0.67 for VLDP (Figure 5). This means that the relationships at the markers overpredict the relationships at the causal loci. When the difference in allele frequencies of causal loci between populations was not reflected by the markers, regression coefficients of between-population relationships were ∼0.30 (Figure 6). Thus the overprediction of between-population relationships using markers was much larger when the difference in allele frequency of the causal loci between the populations was not reflected by the markers.

The correlation between the relationships at the causal loci and at the markers, i.e., the accuracy of the marker-based relationships, decreased when the density of the markers decreased (Figure 5 and 6). When the difference in allele frequencies of causal loci between populations was reflected by the markers, the correlation for within-population relationships was ∼0.91 for HDP and LDP, and ∼0.88 for VLDP. The correlation for between-population relationships was ∼0.70 for HDP and LDP, and 0.60 for VLDP. The correlation between relationships at causal loci and at markers was much lower when the difference in allele frequencies of causal loci between populations was not reflected by the markers (within-population relationships: ∼0.66 for HDP and LDP, ∼0.63 for VLDP; between-population relationships: ∼0.09 for HDP and LDP, ∼0.08 for VLDP).

## DISCUSSION

The objective of this study was to investigate whether differences in LD and allele frequencies of markers and causal loci between populations affect bias of the estimated genetic correlation between populations. Results showed that when a difference in allele frequencies of causal loci between populations was reflected by the markers, estimated genetic correlations were only slightly underestimated using markers. This was even the case when LD patterns, as measured by LD-statistic *r*, were different between populations. When the difference in allele frequencies of causal loci between populations was not reflected by the markers, genetic correlations were severely underestimated. Differences in LD and allele frequencies of causal loci between populations only had a very slight effect on the precision of the estimated genetic correlation.

### Estimating the genetic correlation using marker-based relationships

Genetic variance and heritability estimates are known to be biased when the regression coefficient of the true relationships on the marker-based relationships is not equal to one, i.e., when *E*(**G**_causal loci_|**G**_markers_) ≠ **G**_markers_ (Yang *et al.* 2010; Goddard *et al.* 2011; Yang *et al.* 2015). When this regression coefficient is below one, relationships at the markers show too much variation, resulting in an underestimation of the genetic variance. Yang *et al.* (2010) argued that a regression coefficient smaller than one can be a result of two effects; 1) sampling error on the relationships because the number of markers is finite, and 2) a difference in allele frequency distribution between causal loci and markers. In all our scenarios, the number of markers was finite and the allele frequency distribution was different for causal loci than for markers. However, within populations, the estimated genomic heritability (De los Campos *et al.* 2015) was close to the simulated trait heritability for all scenarios. This suggests that enough markers were used to constrain the sampling error on within-population relationships to an acceptable level, and that our estimated genetic variances were only slightly affected by the difference in allele frequency distribution between causal loci and markers. Thus the underestimation of the genetic correlation between populations is not a result of biased genetic variance estimates.

The relative sampling error as a result of using a finite number of markers was much larger for between-population relationships than for within-population relationships, because more markers are needed to accurately estimate the small between-population relationships (Goddard *et al.* 2011). Moreover, the accuracy of predicting the between-population relationships at the causal loci using markers was depending on the reflection of the difference in allele frequency of causal loci between populations by the markers. Those two effects can result in an underestimated genetic covariance between populations, which can explain the slight underestimation of the genetic correlation in the scenarios where the difference in allele frequencies of causal loci between the populations was reflected by the markers, and the more severe underestimation in the scenarios where this was not the case. The higher sampling error on between-population relationships can also explain the larger underestimation of the genetic correlation for VLDP markers than for HDP and LDP markers. Thus for estimating the genetic correlation between populations, it is important that the difference in allele frequencies of causal loci between the populations is reflected by the markers and that the number of markers is high.

### Regression of the maker-based relationships

Regressing **G** towards the pedigree relationship matrix is a way to correct the marker-based relationships for the sampling error as a result of using a finite number of markers (Powell *et al.* 2010). The regression was strongest for VLDP markers, where it reduced the underestimation of the genetic correlation. Those results agree with the findings that regressing **G** is important when the number of markers is low (Yang *et al.* 2010) and supports our statement that relationships at VLDP markers were affected by sampling error. However, regressing **G** increased the underestimation of the genetic correlation with HDP and LDP markers. The reason for this is not clear. It might be that the regression of **G** not only reduces the sampling error, but also amplifies the effect of the difference in allele frequency distribution of causal loci and markers.

In our study, regressing **G** towards **A** was detrimental for estimating the genetic correlation when using HDP (200 000) or LDP (20 000) markers, where all regression coefficients were close to one, and regressing was beneficial when using VLDP (2000) markers, where regression coefficients were considerably below one. The simulated genome was about one third of the genome of livestock species such as cattle and chicken (Ihara *et al.* 2004; Groenen *et al.* 2009). This would indicate that regressing **G** is detrimental when using a genome-wide total of 60 000 or more markers in livestock. Note that this number of markers will depend on the consistency in LD between populations. Between-population relationships are all closer to zero when consistency in LD between populations is lower (Goddard 2009). Those lower relationships generally require more markers to reduce their relative sampling error to an acceptable level (Yang *et al.* 2010). Hence, we think that the regression coefficients may be a better indicator for deciding whether or not to regress **G**; when all regression coefficients are close to one, e.g., above 0.95, it is probably better to not regress **G** towards **A** when estimating the genetic correlation between populations.

The coefficients to regress **G** towards **A** were approximated using the number of markers and the variation in **G**_markers_**-A**, assuming that the sampling error was only a result of using a limited number of markers (Goddard *et al.* 2011). To investigate the impact of this approximation and whether we could remove the observed underestimation of the genetic correlation by rescaling **G**_markers_ such that *E*(**G**_causal loci_|**G**_markers_) = **G**_markers_, we repeated some analysis using 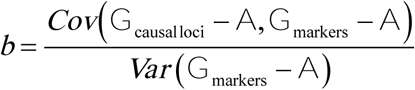 (Goddard *et al.* 2011) as regression coefficient to regress **G** towards **A**. This regression requires the causal loci to be known, which was the case in our simulations. We calculated *b* separately for within- and between-population relationships, using 11 bins based on pedigree relationships within populations (<0.05, 0.05-0.10, 0.10-0.15, 0.15-0.20, 0.20-0.25, 0.25-0.30, 0.30-0.35, 0.35-0.40, 0.40-0.50, >0.50, self-relationships) and 3 bins based on genomic relationships between populations (<0.10, −0.10-0.10, >0.10), and used those *b*’s to rescale the relationships. As shown in Figure 7, this rescaling almost completely removed the bias in genetic correlation estimates using HDP and LDP markers. The genetic correlation was overestimated when using rescaled relationships based on VLDP markers. This might be a result of the much larger sampling error for VLDP markers compared to HDP and LDP markers, which could result in underestimated *b* values. Thus, there appears to be a lower boundary for the number of markers to calculate between-population genomic relationships that can be corrected using regression. Altogether, those results confirm that for an unbiased estimate of the genetic correlation between populations, the regression coefficient of true relationships on marker-based relationships should be one.

**Figure 7.**
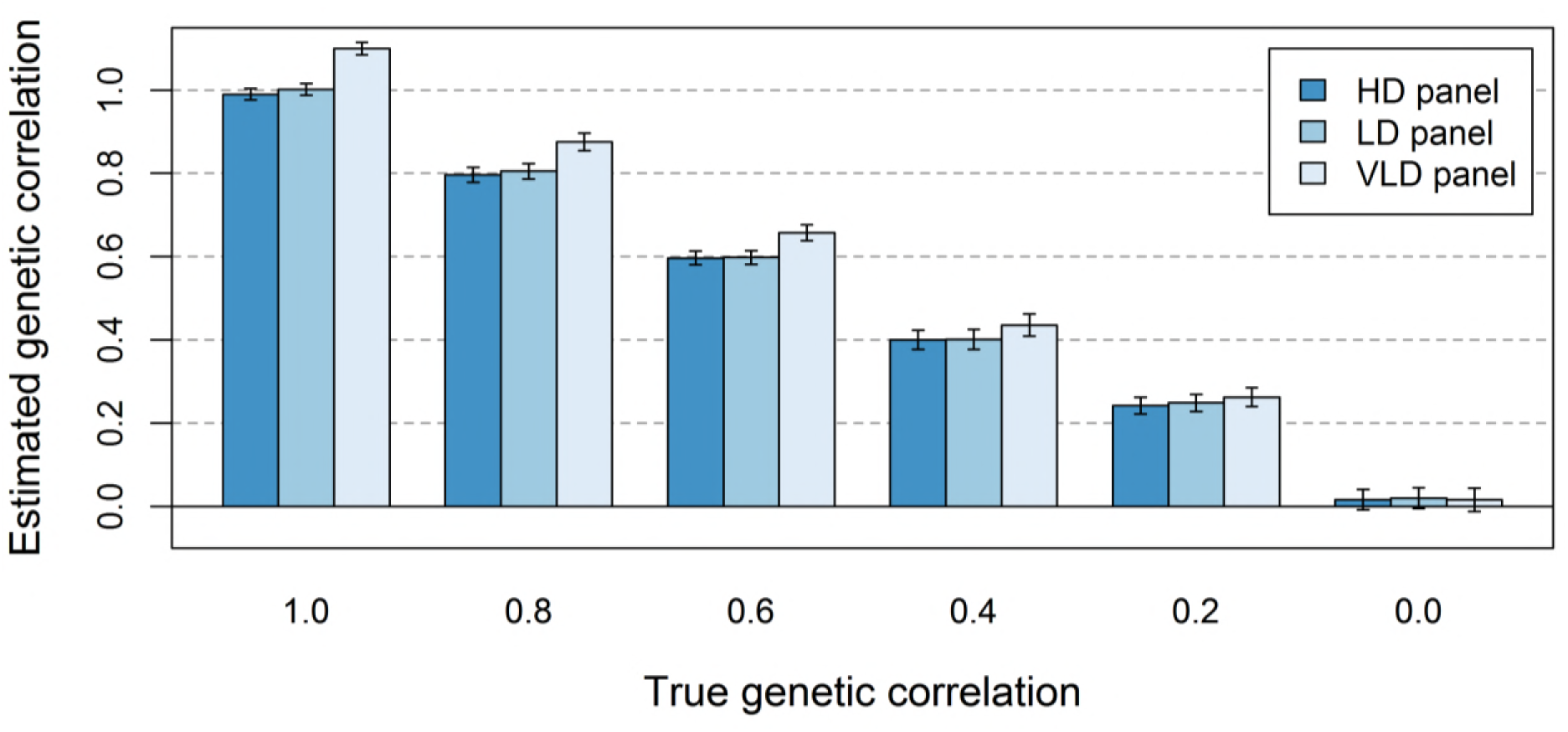
- Estimated genetic correlations between populations after rescaling the marker-based genomic relationship matrix. The average estimated genetic correlation (± standard error) at different simulated genetic correlations for the scenario where markers and causal loci have similar allele frequencies in the two populations when the genomic relationship matrix is either based on the genotypes of HDP (200 000), LDP (20 000), or VLDP (2000) markers, after rescaling the marker-based relationships using a regression coefficient based on the relationships at causal loci. Standard errors were calculated as the standard deviation over replicates divided by the square root of the number of replicates.

### Consistency in LD

We used different marker densities to represent differences in consistency in LD between populations. We expected that a lower consistency in LD would reduce the estimated genetic correlation between populations, because it reduces the correlation between (apparent) marker effects. Surprisingly, our results showed that estimated genetic correlations were similar with HDP and LDP markers, and only slightly lower with VLDP markers. This can be explained by the potential of marker-based relationships to accurately predict the relationships at the causal loci, which is essential to unbiasedly estimate the genetic (co)variances and the genetic correlation between populations. A lower consistency in LD between populations results in a lower variation in between-population relationships (Goddard 2009; Goddard *et al.* 2011). Because a lower consistency in LD reduces the variation in between-population relationships at both causal loci and markers, the regression coefficient of the relationships at the causal loci on the relationships at the markers may not be affected much (Figure 5 and 6; HDP and LDP markers). Therefore, the estimated genetic correlation between populations seems little affected by the consistency in LD between the populations.

The consistency in LD between populations does affect the correlation between the relationships at the causal loci and the marker-based relationships (Figure 5 and 6), i.e., the accuracy of the marker-based relationships. For an unbiased estimate of the genetic correlation between populations, the regression of true relationships on marker-relationships should be one and marker-based relationships don’t necessarily have to be accurate. This is in contrast to estimating genetic values, as is done in genomic prediction, for which relationships have to be accurate and have to show variation (Goddard *et al.* 2011). Thus, an unbiased estimate of the genetic correlation between populations does not guarantee that accurate genomic prediction across populations can be performed.

### LD structure

The extent and consistency of LD in the simulated populations is comparable to the patterns found in chicken and pig populations (Andreescu *et al.* 2007; Badke *et al.* 2012; Veroneze *et al.* 2013; Veroneze *et al.* 2014). This simulated LD was much higher than generally found in human populations (Pritchard and Przeworski 2001; Shifman *et al.* 2003). Since marker density, and thereby the average LD between causal loci and nearest marker, had no effect on the estimated genetic correlation, it is expected that the simulated LD pattern did not affect the results.

We simulated causal loci randomly spread across the genome, which is not always the case in real populations. When causal loci are enriched in regions with either high or low LD, (co)variance estimates can be over- or underestimated (Speed *et al.* 2012; Yang *et al.* 2015). However, we would expect a smaller impact of the heterogeneity of LD on the estimated genetic correlation than on the heritability, since differences in LD across the genome affect both the genetic variance and covariance estimates. This mechanism may also explain why genetic correlation estimates between traits within a population are less affected by incomplete LD between causal loci and markers than genetic variance estimates (Trzaskowski *et al.* 2013).

### Genomic relationship matrix

The current generation within each population was used as base population for our genomic relationships, since we used current population-specific allele frequencies. This means that between-population relationships are on average zero. When the consistency in LD between the populations is not zero, due to the existence of a recent or distant common ancestor, between-population relationships will show variation around zero (Goddard 2009). That variation is essential in order to estimate the genetic correlation between populations, and genetic correlation estimates are more precise when the variation in between-population relationships is higher (Visscher *et al.* 2014).

Another commonly used multi-population **G** matrix is the matrix following Chen *et al.* (2013). We repeated part of our analyses using that matrix, where the scaling factor of the block between populations is 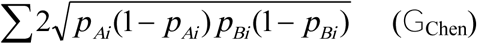 instead of 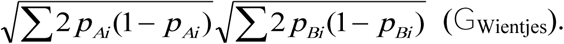 In agreement with our previous study based on causal loci (Wientjes *et al.* 2017), we found that genetic correlations were underestimated using **G**_Chen_. This underestimation is mainly a result of effectively removing markers segregating in only one population from the scaling factor of between-population relationships. This underestimation increases when those markers were also removed from within-population relationships, because it increased the bias in genetic variance estimates. Moreover, **G**_Chen_ was more prone to singularities than **G**_Wientjes_. In **G**_Wientjes_, markers segregating in only one population contributed to the scaling factor for between-population relationships, which resulted in lower between-population relationships when the number of markers segregating in only one population was higher. This resulted in a larger difference between within- and between-population relationships in **G**_Wientjes_, which reduced the risk of singularities.

### Implications

Marker panels are generally composed to have intermediate allele frequencies across multiple populations (Matsuzaki *et al.* 2004; Matukumalli *et al.* 2009; Groenen *et al.* 2011). Therefore, markers tend to have a higher average minor allele frequency than causal loci (Yang *et al.* 2010; Kemper and Goddard 2012). Moreover, the difference in allele frequencies of causal loci between populations is probably not accurately represented by markers. Those factors likely result in underestimated genetic correlations between populations using real data, but the impact of each of the factors requires further research.

### Conclusion

For an unbiased estimate of the genetic correlation between populations from marker information, it is important that marker-based relationships accurately predict the relationships at causal loci, i.e., *E*(**G**_causal loci_|**G**_markers_) = **G**_markers_. To achieve this, the difference in allele frequencies of causal loci between the populations should be reflected by the markers, and the number of markers should be sufficiently high to constrain the sampling error on between-population relationships to an acceptable level. The consistency in LD between populations has little effect on the bias of the estimated genetic correlation.

## ACKNOWLEDGMENTS

This study was financially supported by NWO-TTW and the Breed4Food partners Cobb Europe, CRV, Hendrix Genetics and Topigs Norsvin. The use of the HPC cluster has been made possible by CAT-AgroFood (Shared Research Facilities Wageningen UR).

